# High-resolution 3-dimensional micro-CT imaging of the nucleus pulposus using a novel contrast agent

**DOI:** 10.1101/2024.11.25.625182

**Authors:** Madison M Buckles, Nada H. Warda, Jacob C. Moncher, Mohammad Yunus Ansari

## Abstract

**Objective:** The nucleus pulpous (NP) in the intervertebral disc (IVD) is the first structure to exhibit degenerative changes during IVD degeneration (IDD). Currently, micro-computed tomography (micro-CT) imaging of the NP is a limiting factor in detecting IDD at an early stage. In this study, we used potassium iodide (KI) as a contrast agent to stain the NP in the IVDs of the tails and spines of mice and rats and imaged them with a micro-CT scanner.

**Methods:** We collected tails and spines from 12-week-old mice and rats and stained them with KI followed by micro-CT imaging. To induce IDD, we performed caudal disc needle puncture surgery (NPS) in age and sex-matched mice (n=5) and stained the tail with KI for imaging with micro-CT to study disc degeneration.

**Results:** KI successfully stained and helped visualize the 3D structure of the NP, with X-ray attenuation properties comparable to bone. Contrast staining with KI enabled accurate and reproducible quantification of disc height and NP volume. The cross-sectional micro-CT images of NPS discs were indistinguishable from the histological findings of the same sample and showed similar degenerative changes in the NP. Additionally, we estimated that up to 15 sagittal sections, each 5μm thick with 75μm spacing, would be needed to fully assess disc degeneration.

**Conclusion:** This study demonstrated that KI can be used to positively stain NP in the intact tail or spine and provide qualitative and quantitative data without any adverse effect on the histological processing of the samples.

## Introduction

The intervertebral disc (IVD) is a fibrocartilaginous structure between two adjacent vertebrae in the vertebral column, providing flexibility and dampening loads in the spine. The IVD consists of a central, proteoglycan-rich, jelly-like nucleus pulposus (NP) surrounded by lamellae of annulus fibrosus (AF), which is sandwiched between two vertebrae with cartilaginous end plate (Supplementary Figure 1) (1). The NP tissue plays a vital role in the function of the IVD, and changes in NP structure promote IVD degeneration and low back pain (2).

Intervertebral disc degeneration (IDD) is highly prevalent and a leading cause of low back pain (LBP), representing a significant socioeconomic burden costing billions of dollars in healthcare costs and lost productivity (3, 4). A better understanding of the mechanisms and structural and molecular changes in IVD during IDD is vital for the development of therapeutic strategies for the treatment of low back pain. One of the challenges associated with understanding the mechanisms of IDD is the lack of high-resolution 3D imaging techniques to visualize the NP. Imaging by micro-computed tomography (micro-CT) relies on the attenuation of X-rays passing through tissues up to the micrometer resolution limit, however, soft tissues such as IVDs have low X-ray attenuation properties and are difficult to visualize without the help of a contrasting agent. Contrast-enhanced micro-CT is an important tool for studying the three-dimensional microstructures of soft tissue, which otherwise have low X-ray attenuation properties (5).

Small animal models, such as mice and rats are of paramount importance for studying the mechanisms of disease pathogenesis due to the availability of genetically engineered animals and the feasibility of the work. However, current micro-CT imaging to examine changes in the IVD is based on a negative staining approach where the NP is not visible, and changes in disc height and other bone parameters are considered as damage to the NP and overall disc structure (6). Many studies have shown that contrast-enhancing agents such as Lugol’s solution (Iodine-potassium iodide), Hexabrix (ioxaglate), Cysto Conray II (iothalamate), CA^4+^, PMA (phosphomolybdic acid), and PTA (phosphotungstic acid) improve the visibility of soft tissues by micro-CT (7, 8). However, these approaches take a long time and preferentially stain the AF. In this study, we demonstrate a quick, positive staining approach of the NP by potassium iodide (KI), resulting in the visualization of the NP by micro-CT. We show here that KI differentially stained the NP tissue in mouse and rat IVDs and enabled us to image the 3D structure of the NP at a 10μm spatial resolution. Moreover, with this new method, we can quantify the volume of both a healthy and a degenerated NP in a mouse model. Here, we show that KI can be used as a contrast agent to visualize the NP.

## Materials and Methods

### KI staining and micro-CT of mouse tails

All the animal studies were approved by the Institutional Animal Care and Use Committee (IACUC) of Northeast Ohio Medical University. C57BL/6J mice were obtained from Jackson’s Laboratory (stock number # 000664) and housed in standard cages with ad libitum access to water and food. Male and female mice (n=3 to 5 per experimental group) were euthanized at 12 weeks to collect the spines and tails for KI staining. Skin-free tails were fixed in 10% neutral buffered formalin for 24 hrs at room temperature, washed three times in PBS, and then placed in KI staining solution for the allotted time (30 mins, 2 hrs, 4 hrs). KI was purchased from Fisher Scientific (#BP367), and a 50% KI stock solution in water was prepared, which was further diluted to obtain 25% and 12.5% KI using mQ water. All solutions were stored at RT in a dark environment. A SkyScan 1273 micro-CT scanner (Bruker) was used to assess the IVD morphology using a 10-μm isotropic voxel size. Scanner settings were maintained at 90 kV of tube voltage and 166 μA of tube current through a 0.5mm Al filter. Scanned projections were reconstructed in NRecon (Bruker) and analyzed using Avizo 3D Pro (ThermoFisher). Percent X-ray attenuation was determined using ImageJ.

### Disc needle puncture surgery (NPS) induced intervertebral disc degeneration (IDD) in mice

IDD in mouse caudal discs was established by annular puncture using a 30-gauge needle, as described in (9). Briefly, 12-week-old C57BL/6J male and female mice (n=5 per group) were anesthetized by exposure to isoflurane and given buprenorphine for pain relief. The discs were located by palpation, and a 5mm longitudinal incision was made on the skin to expose the disc. The caudal IVDs (Ca4/5 and Ca6/7) were punctured using a 30-gauge needle at a controlled depth of needle bevel under a stereoscope. No leakage of disc tissue was observed after the NPS. The skin was then closed using tissue glue. Two discs per mouse were punctured with adjacent discs used as sham controls. The mice were euthanized 2 weeks post NPS, and the tail samples were collected, fixed in 10% NBF, and stained with KI for micro-CT scanning.

### Histology

Reversal of the KI stain was performed through alcohol leeching. The tail samples were processed for histology as described previously (10). Briefly, the mouse tails were dehydrated with a series of ethanol and xylene washes followed by embedding in paraffin. The tissue was cut into 6μm sagittal sections and stained with Hematoxylin & Eosin (H&E) for imaging. The images were captured using a slide scanner (#Olympus VS120). The NPS discs were scored according to the MERCY scoring system to determine the degree of disc degeneration (11).

### Statistical analysis

The graphical data is expressed as mean±SD. Statistical analysis was conducted using GraphPad Prism software (Version 10). The data was analyzed using Student’s t-test. *P* < 0.05 was considered statistically significant.

## Results

### Optimization of IVD staining with KI

The AF, NP, and cartilage endplate tissues of IVDs do not attenuate the X-rays and thus are invisible in micro-CT scanning experiments (Figure 1A). To study the structure of IVD tissues, staining of tissues with contrast-enhancing agents is a standard practice to visualize tissues that normally cannot be visualized by micro-CT. To visualize the IVD tissues, mouse tails ranging from Ca3 to Ca9 were stained with different concentrations of KI (50%, 25%, and 12.5% w/v) for the indicated time (0.5 hours, 2 hours, and 4 hours) followed by scanning by micro-CT machine. KI fully penetrated the disc and provided a clear radiographic contrast to NP as early as half an hour with almost every concentration tested in this study (Figure 1B). The NP and AF tissues were not visible using the same micro-CT scanning parameters (Figure 1A), showing the difference between stained and unstained IVDs. We observed differential staining of AF and NP in KI-stained discs with high KI uptake in the NP compared to AF. There was no macroscopic structural change observed in the tail after staining with KI (Figure 1D). In our staining optimization, we found that 25% KI in 0.5 hours resulted in the best staining and radiographic contrast of mouse tail NP tissue as shown by attenuation intensity profile data (Figure 1C), and was used in further staining studies. These results show a positive staining of NP tissue in a healthy IVD.

**Figure 1:**
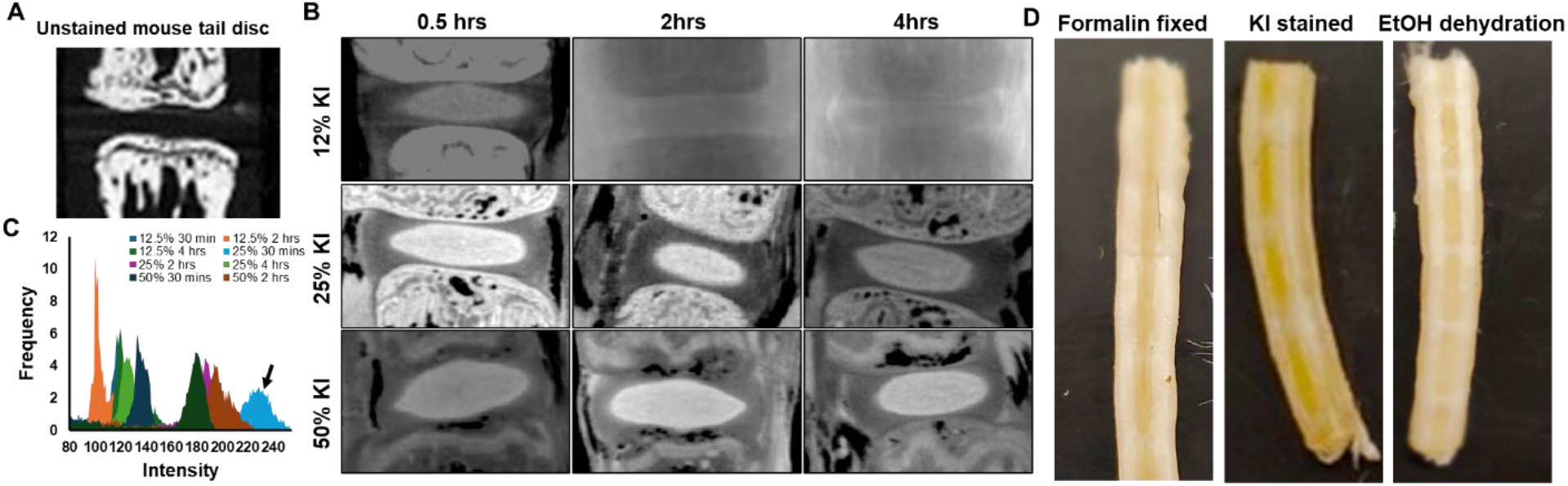
Optimization of KI staining of mouse tail IVDs. (A) Representative micro-CT image of an unstained mouse tail IVD. (B) Mouse tails were collected from 12-week-old C57BL6 male and female mice (n=3 to 5 per concentration, per time point) and fixed in 10% NBF for 24 hours followed by staining with different concentrations of KI for specified time. The samples were scanned with micro-CT as described in the materials and methods. (C) KI staining intensity of NP was estimated by ImageJ. 25% KI for 30 minutes showed maximum attenuation intensity (indicated by black arrow). (D) Representative images of formalin fixed, KI stained, and EtOH dehydrated mouse tail samples showed no macroscopic damage to the tissue.

### Disc degeneration analysis by KI staining in an NPS model

Previous studies using an iodine-based contrast agent (Hexabrix) to visualize discs have shown a negative correlation of NP staining with healthy NP stains less, and damaged NP allows high penetration with the contrasting agent (7). We tested our KI-based positive staining method to determine the damage to NP in an NPS model of disc degeneration. The NPS was performed by puncturing the caudal disc using a 30-gauge syringe needle as shown in the flow chart (Supplementary Figure 2). KI staining showed a decrease in NP area in the NPS disc, compared to adjacent sham control discs (Figure 2A). The 3D rendering of the disc clearly showed damage to the NP in the NPS model (Figure 2B). We also determined the disc height (Figure 2C) and NP volume (Figure 2D) which showed a significant reduction in NPS disc compared to sham. The observations in contrast-enhanced micro-CT data were tested by histology using H&E staining and displayed similar results between histology and micro-CT (Figure 2E). We also scored the damage to NP (Figure 2F) using the MERCY scoring system (11). These results show that KI can be used as a contrast agent to visualize damage to NP. In addition to providing contrast to NP, KI also helped visualize the osteophytes which can be seen in 3D rendered images (Figure 2B).

**Figure 2:**
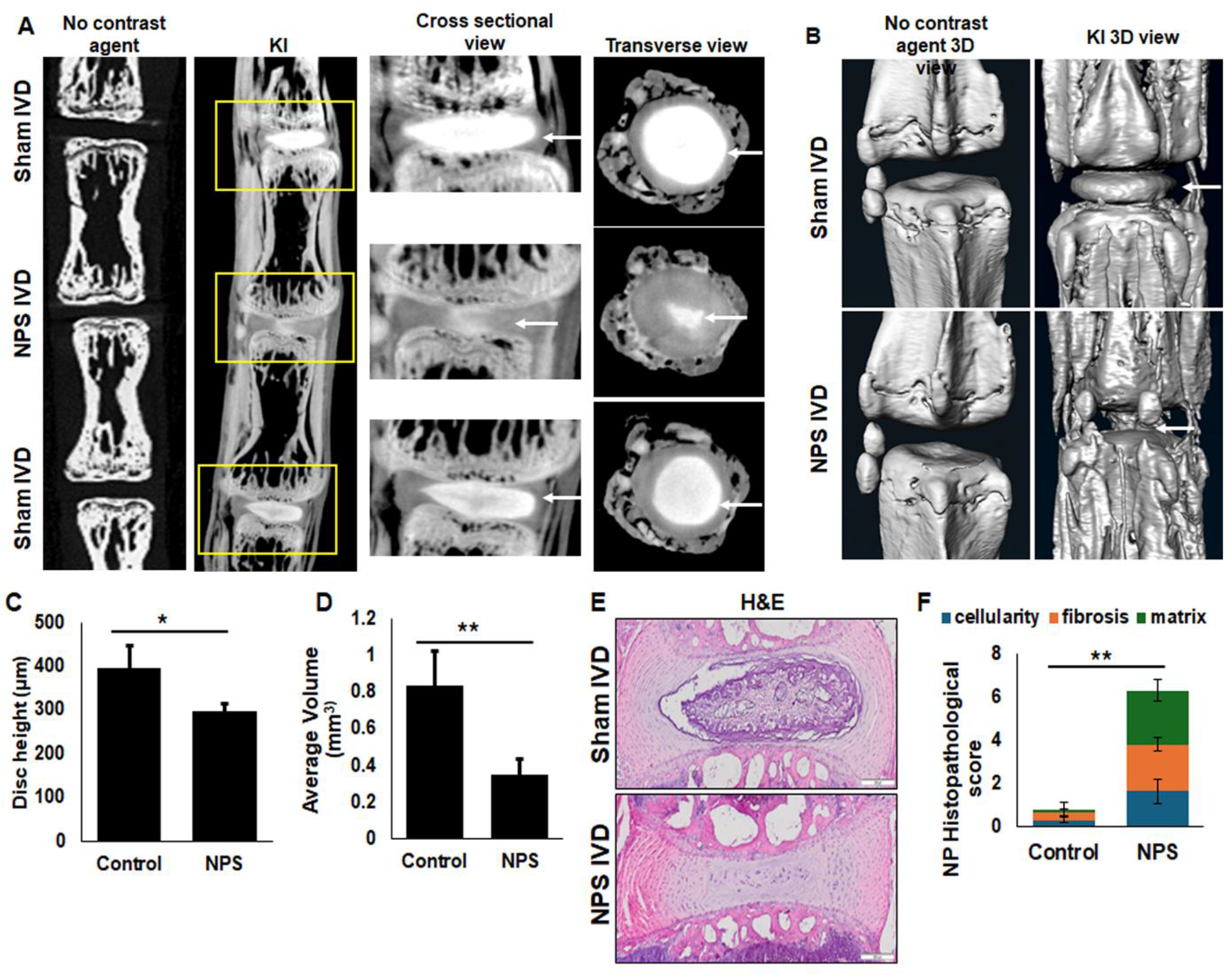
Visualization of damage to the NP in a needle puncture surgery (NPS) model of disc degeneration in mice. (A) 12-week-old C57BL/6J mice (n=5) were subjected to NPS. The tail samples were collected 2 weeks post-NPS and processed for KI staining and micro-CT imaging. Representative micro-CT images of NPS and sham IVDs from unstained (no contrast agent) and KI-stained mouse tails. The NP is highlighted by the white arrow in the Cross-sectional and transverse zoomed view. (B) Representative 3D view of the sham and NPS IVDs showed a drastic difference in the shape and appearance of NP upon needle puncture. White arrows indicate the NP. (C) The disc height in the KI-stained control and NPS disc showed a significant difference (Student t-test, * p<0.05). (D) The volume of NP in the sham and NPS IVDs showed a significant loss of NP after needle puncture (Student’s t-test, * * p<0.005). (E) The tails were processed for histology and stained with H&E. Representative H&E-stained images of control and NPS IVDs. (F) MERCY score of NP in the mouse tail showed significant IVD degeneration. (Student t-test, p<0.05). * * p=0.04 for cellularity, p=0.01 for fibrosis, and p=0.002 for matrix. These results showed that KI staining can visualize NP damage.

### Visualization of mouse spine disc with KI staining

To test KI staining for spine IVDs, we stained the mouse lumbar spine with 25% KI solution initially for 30 minutes, but the micro-CT images were not as sharp as the tail images (data not shown). We then increased the staining time from 30 minutes to 45 minutes. As shown in Figure 3A, the NP tissue in the spine disc was visualized by increasing the time of incubation with KI. These micro-CT images of spine discs (cross-sectional and 3D view) show that KI works well with IVDs from the tail and the spine.

**Figure 3:**
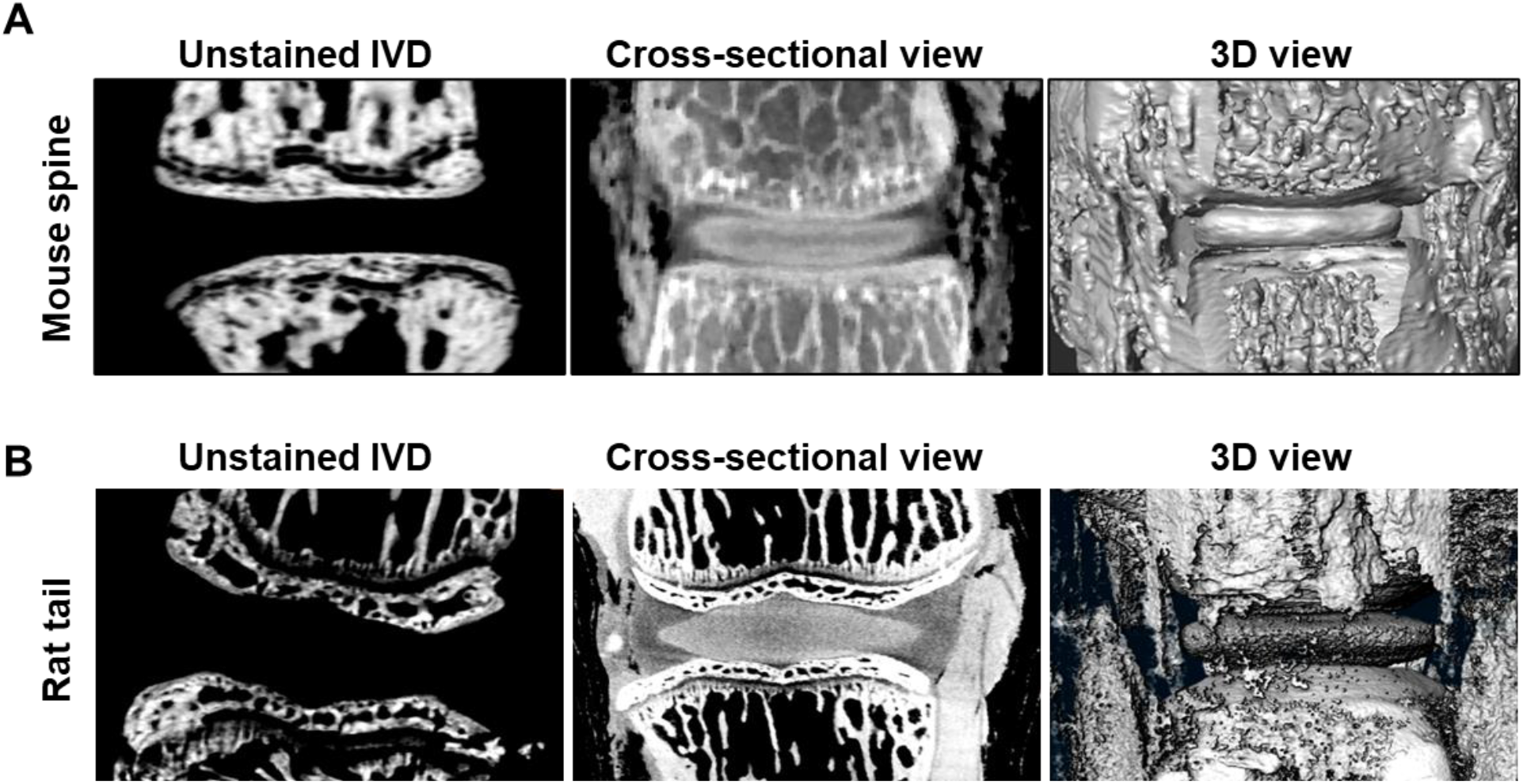
Visualization of NP in mouse spine and rat tails with KI staining. (A) The spine was collected from mice (n=3) and incubated in KI solution for 45 minutes followed by scanning with a micro-CT machine. The unstained spine was scanned as a control. (B) The rat tail (n=3) was incubated in KI solution as above and scanned with a micro-CT machine. The images show the staining of NP with KI in the cross-sectional and the 3D-rendered view.

### Visualization of rat tail disc with KI staining

We further tested KI to visualize NP in the IVDs of larger rodents. We stained rat tails with KI for 45 minutes and scanned the disc with micro-CT. Interestingly, KI was found to penetrate the disc in rat tail resulting in the visualization of the NP tissue (Figure 3B). These results show that KI can be used as a contrast agent to enhance the visualization of the NP by micro-CT.

## Discussion

The current histological approach of evaluating changes in NP in the degenerating disc is time-consuming, destructive and cannot asses the 3D spatial morphology of the NP. In addition, the semi-quantitative scores are subjective with intra- and inter-observer variations. The 3D morphometric analyses obtained through micro-CT offer a distinct advantage over the 2D histological approach by providing accurate volumetric assessment and eliminating the need for sample positioning, and possible artifacts introduced by dehydration, embedding, and sectioning. Moreover, the samples can still be processed for histology after micro-CT.

Contrast-enhanced imaging of soft tissues, which otherwise have low to no X-ray attenuation properties, is the method of choice to visualize them by CT scans. Iodine-based contrast agents such as Lugol’s solution, Hexabrix, and Cysto Conray II, are commonly used due to their high X-ray attenuation properties (7, 8). Here, we show that KI can be used as a contrast agent to visualize NP tissue in the IVD. The NP is a water-rich, gelatinous structure with a high level of glycosaminoglycans which gives it an overall negative charge. It has been reported that the Hexabirx staining of NP tissue is inversely proportional to the amount of GAG in the NP due to its negative charge (7). Previous studies provide a negative approach to visualizing changes in NP that rely on not staining the NP. Our method provides a positive staining of NP in which we stain the NP to visualize through micro-CT and we can directly measure the changes in the NP.

This positive staining of NP in the IVD provides us with a huge opportunity to study IDD in animal models. This method can also be optimized for other animal models and to study IDD without the dehydration of the disc and without going for a longer incubation period (2 months incubation in (12)). Other published studies require cutting and isolation of the IVD from the vertebral column or tail vertebrae (12) which poses additional challenges and may require expertise and special skills and additional tools such as the LASER dissection tool for precise cutting to prevent any damage to the IVD. The above study also showed major macroscopic structural changes including damage to NP and separation of AF lamellae in the isolated disc due to fixation and longer staining time while we did not see any macroscopic changes in the IVDs (Figure 1D) and processed the tissue for histology and did not see any adverse effect of KI on disc histologically. In this method, the NP tissue was stained while present in the spine or tail giving an added advantage to analyzing the bone in the same scan. The other advantage of this method is we get to see the whole NP tissue in a 3D view which helps in the analyses of the entire NP in the disc rather than looking at thin sections by histology which shows only a small portion of the NP. The staining of KI is reversible giving another advantage of processing the disc samples for further histological analysis. In addition to the NP, other soft tissues around the disc were also visualized by KI staining and can be studied. KI can also be used for IVD characterization in genetically modified mice and rats to explore any phenotype. This method can also be used to assess the health and regeneration of NP in cell transplantation therapy. The X-ray attenuation by 25% KI-stained NP was as high as by bone suggesting that the imaging can be performed at lower voltage or mAmp under clinical settings to minimize radiation exposure while maintaining the signal-to-noise ratio (13).

The other implication of our study is the clinical diagnosis of disc degeneration. Currently, iodine-based contrast agents such as ioxaglate are used widely for CT scans for the diagnosis of various illnesses (14). Due to quick staining time, and the ability of KI to stain the NP at the lowest concentration (12%) tested in this study, this method can be developed for use in X-ray CT scans for IDD in patients.

The limitations of this study include the lack of visualization of AF lamellar structures, however, I2KI staining is known to have limited revealing of tissue sub-structures and is more suitable for larger tissue or organs in which overall anatomical structure is the subject of study (15). PTA and PMA were shown to stain collagen-rich AF lamellae better than I2KI (12).

In summary, this is the first study to show that the NP tissue in intact IVD can be visualized by KI staining within 30-45 minutes. We showed here that contrast-enhanced micro-CT can be used to analyze and quantify changes in NP structure in animal models of IDD and propose that this method can be used to study changes in IVD with age or other conditions. This is significant as the mechanism of disc degeneration is not fully understood, and efficient assessment of NP degeneration is required for the successful treatment of IDD. This study provides a platform to evaluate disease-modifying drugs for the treatment of IDD. This approach has the potential to become a standard analysis method for comprehensive analyses of disc degeneration and treatment efficacy.

## Acknowledgements

We thank Sharon Usip and David Waugh for their help with the micro-CT scanner. We thank Dr. Nic Leipzig, University of Akron for rat tails. We thank NEOMED for funding to support this project.

