## Supplementary data for "High-resolution 3-dimensional micro-CT imaging of the nucleus pulposus using a novel contrast agent"

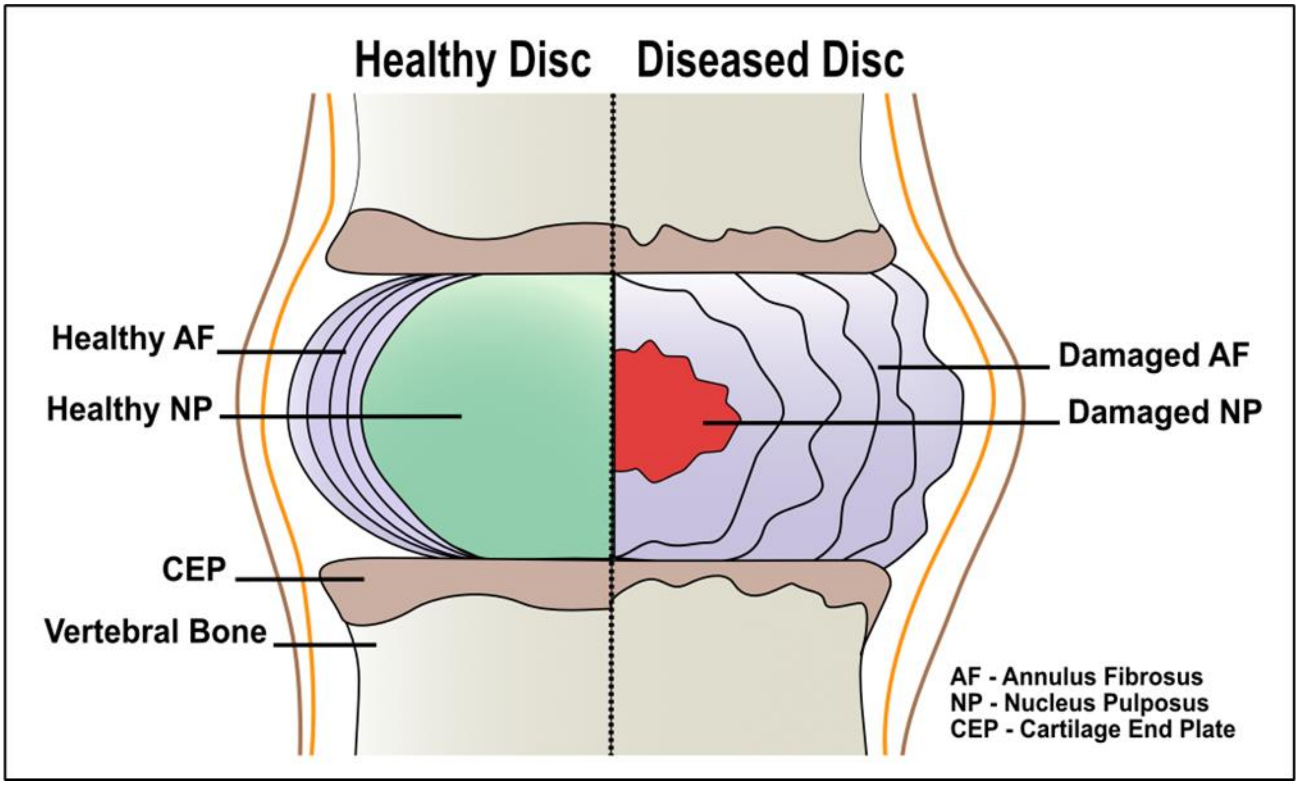


**Supplementary Figure 1:** Schematic representation of a healthy and degenerated intervertebral disc (IVD) structural organization. The degenerative changes in the IVD include loss of nucleus pulposus (NP), tear in annulus fibrosus (AF) lamellae, and damage to the cartilaginous end plate (CEP).

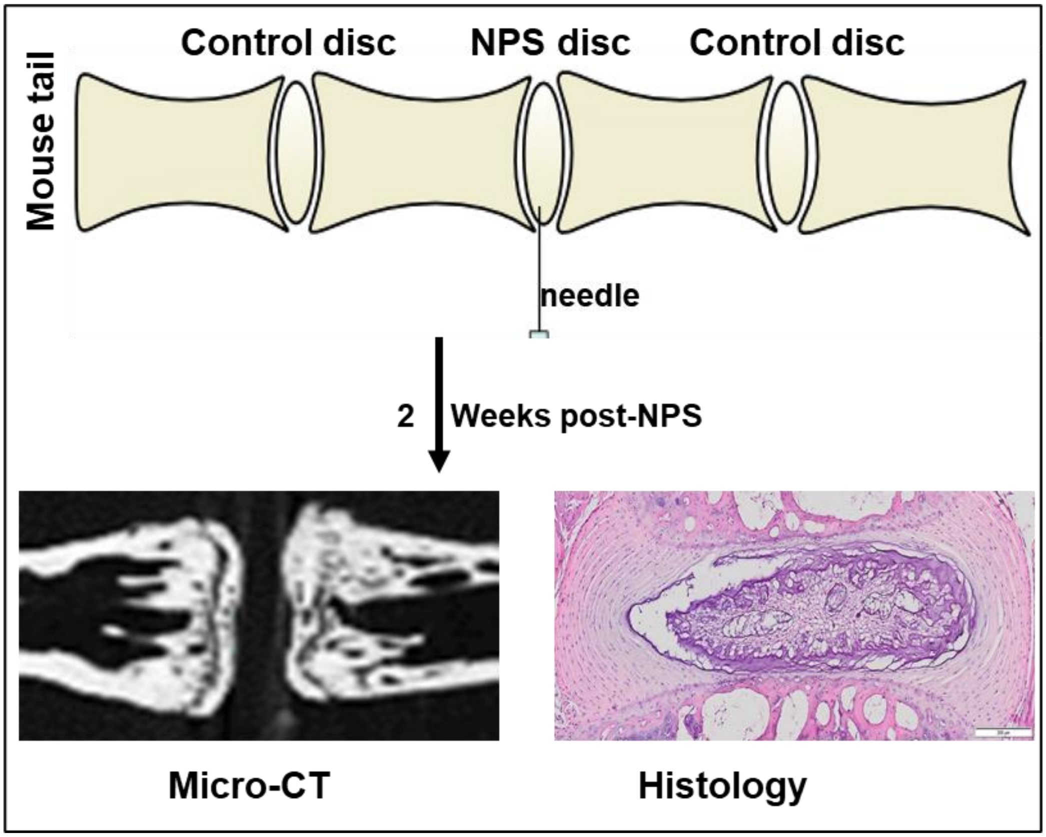


**Supplementary Figure 2:** Flow chart of the study (validation or testing of the KI staining method to determine disc degeneration in needle puncture surgery (NPS) model). Briefly, the mouse tail disc was punctured using a 30-gauge needle. The mice were sacrificed 2 weeks post-surgery and the IVD was analyzed by microCT and histology.
